# Transarterial embolization modulates the immune response within target and non-target hepatocellular carcinomas

**DOI:** 10.1101/2020.11.07.372896

**Authors:** David Tischfield, Alexey Gurevich, Omar Johnson, Isabela Gayatman, Gregory J. Nadolski, David E. Kaplan, Emma Furth, Stephen J. Hunt, Terence P. F. Gade

## Abstract

Recent successes in the field of immuno-oncology have generated considerable interest in the investigation of approaches that combine standard of care treatments with immunotherapies. Transarterial embolization (TAE) represents an attractive candidate for this approach given the potential for immune system stimulation, however little is known about the influence of TAE on the tumor immunoenvironment. The purpose of this study was to perform a cellular analysis of tumor infiltrating lymphocytes (TILs) and PD-L1 expression following TAE in a translational rat model of hepatocellular carcinoma (HCC) and to identify factors that influence this response. We show that TAE causes dynamic changes in immune cell populations, with variable increases in CD3, CD4, and CD8 cells within embolized tumors over time. We also show that TAE alters the immunobiology of distant, non-target tumors as demonstrated by an increased number of CD4, CD8, and Foxp3+ cells within the intratumoral compartment of non-target tumors. We demonstrate that, in response to TAE, tumor cells up-regulate expression of PD-L1. Finally, we demonstrate marked differences in terms of the foreign body reactions induced by two commonly used embolic particles, and show changes in lymphocyte and macrophage recruitment that depend on the type of embolic particles and their propensity to extravasate beyond the vasculature and into the tumor parenchyma. These findings hold important implications for the on-going development of novel therapeutic strategies combining locoregional therapy with immunomodulators, as well as for the development of techniques and materials that can further leverage the unique modulation of the tumor immune microenvironment.

## Introduction

Hepatocellular carcinoma (HCC) is the second-leading cause of cancer-associated death worldwide and the most rapidly increasing cause of cancer death in the United States^1–3^. Nearly 65% of HCC patients are diagnosed at an intermediate or advanced stage, when curative therapies, such as resection or transplantation, are no longer feasible^4^. As a result, locoregional therapies (LRT) including transarterial embolization (TAE) and transarterial chemoembolization (TACE) are the most common treatments for HCC in the United States. TA(C)E exploits the vascular biology of HCC to deprive tumors of essential nutrients through the intra-arterial administration of embolics with or without chemotherapeutics, leading to growth arrest and/or necrosis^5,6^. While TA(C)E has a proven survival benefit, local and distant recurrence are common, and long-term survival rates are poor^7,8^. This limitation has spurred interest in leveraging the unique modulation of the tumor microenvironment enabled by TA(C)E through combination therapy in order to generate a more durable response.

In recent years, immunotherapy has become an established pillar of cancer therapy. While preliminary trials using immune checkpoint inhibitors (ICIs) to treat HCC have shown encouraging results, the majority of patients continue to progress^9,10,11,12,13^ As such, there is growing interest and enthusiasm for the investigation of approaches that combine standard of care treatments, such as embolotherapy, with immunotherapy to overcome these limitations. Multiple LRTs including embolotherapies have shown the potential for immune activation^14,15,16,17,18,19^. TA(C)E represents a particularly attractive candidate for this approach given the potential for immune system stimulation through the large-scale release of tumor-associated antigens (TAA) following ischemia-induced cell death, without the denaturation of these proteins that can accompany thermal ablation strategies^18^.

While the advent and successful application of immunomodulators, including immune checkpoint inhibitors, suggest great promise for synergism in combination with embolotherapy, the realization of this promise has been limited by a dearth of preclinical studies characterizing its influence on the tumor immune micorenvironment^20,21^. Indeed, while clinical trials combining TACE with ICIs are ongoing^22,23^, there has been limited characterization of the immunobiology of embolotherapy that is required before therapeutic immunomodulation can be integrated effectively into current protocols. Fundamental questions remain to be answered in this regard including the ability of embolotherapy to enhance infiltration and activation of the immune cell populations that are required to target HCC cells.

Two major obstacles in the study of embolotherapy-related immune responses include the lack of standardized cross-institutional protocols and limited number of animal models that faithfully recapitulate the hallmark features of HCC and clinical embolotherapy protocols. HCC generally arises on a background of chronic liver disease or cirrhosis, which profoundly alters the immunobiology and architecture of the liver^24,25^. It is therefore imperative that chosen animal models not only recapitulate the immune microenvironment that HCC arises within, but also the changes in scaffolding and vasculature that accompany chronic liver disease. Complicating matters further is the inconsistent use of standardized embolic agents across studies. Depending on their properties, different embolic agents have varying propensities to extravasate outside of the vasculature and elicit inflammatory tissue reactions^26–31^ – properties which may influence the overall immune response to TACE.

Greater knowledge of the mechanisms of immune activation in the setting of embolization is needed in order to design more effective treatment strategies for maximizing the immune response. Herein, we describe the immunomodulatory effects of TAE on autochthonous HCCs induced in a background of cirrhosis in a translational rat model. Using a combination of immunohistochemistry, *in situ* hybridization, and flow cytometry we compare the abundance of CD3, CD4, CD8, Foxp3, and PD-L1-expressing cells between embolized and non-embolized tumors. In addition, we compare differences in the immune cell recruitment associated with two embolic agents commonly used in the clinic. Finally, we demonstrate that similar phenomenon are present following TACE in human samples.

## Results

### Cirrhosis and TAE Modulate Lymphocyte Populations

As a majority of HCCs in the US arise in a background of cirrhosis, we sought to determine how underlying liver disease affects the immune microenvironment and the lymphocyte response to TAE (Figure 1). To do so, we used flow cytometry to compare tissue lymphocyte populations in three cohorts of animals including: 1) naïve, healthy rats following sham TAE, 2) rats with hepatic cirrhosis following sham TAE, and 3) rats with hepatic cirrhosis following TAE with microspheres. We found that relative to naïve rats, there was a significantly greater proportion of CD8+, CD25+/CD4+, and Foxp3+ lymphocytes in the livers (36.1 vs 47.7, P<0.05; 4.6 vs 34.9, P<0.01; 0.8 vs 9.8, P<0.01) and spleens (39.0 vs 50.9, P<0.05, 4.7 vs 22.9, P<0.05; 0.6 vs 3.9, P<0.05) of cirrhotic rats as well as a decrease in the ratio of CD4+ cells (59.6 vs 46.9, P<0.05; 58.0 vs 43.4, P<0.05) (Figures 2a, 2b, and 2c). Interestingly, we found that at 7 days post-embolization, rats treated with TAE demonstrated a significant reduction in the ratio of CD8+ cells within spleen, liver and tumor (50.9 vs 36.7, P<0.01; 47.7 vs 38.4, P<0.05; 49.1 vs 35.5, P<0.05), with a corresponding increase in CD4+ cells (43.4 vs 60.2, P<0.01; 46.9 vs 57.6, P<0.05; 44.5 vs 58.9, P<0.05) that was not significantly different from the proportions of CD8+ and CD4+ cells found in healthy animals (Figure 2a). In addition, rats with hepatic cirrhosis undergoing TAE demonstrated a significant increase in the absolute number of CD25-/CD4+ as proportion of CD3+ cells relative to rats with hepatic cirrhosis undergoing sham embolization within the spleen, liver and tumor (51.9 vs 34.4, P<0.05; 45.3 vs 30.9, P<0.01; 42.5 vs 24.9, P<0.01) (Figure 2c). Taken together, and consistent with clinical findings^32,33,34^, these data indicate that cirrhosis causes global alterations in the proportions of major interstitial lymphocyte populations, and that TAE appears to normalizes these changes through influx of novel effector T cells..

**Figure 1.**
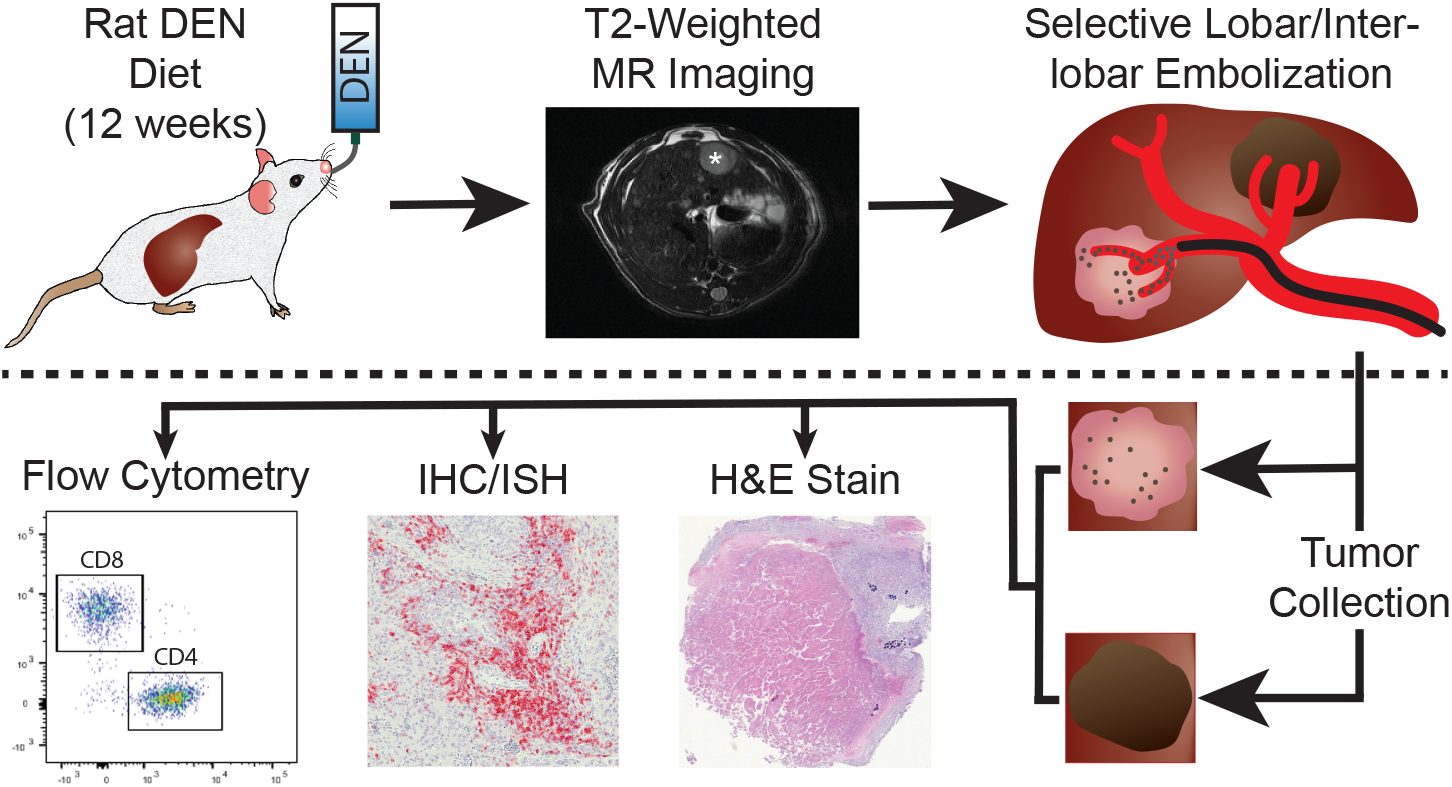
Experimental Schematic. Autochthonous HCCs were induced in male Wistar rats through ad libitum oral administration of 0.01% diethylnitrosamine in their drinking water for a total of 12 weeks. Following completion of the DEN diet, tumor development was monitored using T2-weighted MRI. Tumors reaching 0.5 to 1cm in minimum transverse diameter were selected for experiment and underwent selective transarterial embolization as described previously. At selected post-TAE time points, samples were collected for analysis using either flow cytometry (FC) or immunohistochemistry (IHC)/*in situ* hybridization (ISH). Tumors analyzed using FC were divided into two halves at the time of collection. One half was prepared for histological analysis, including hematoxylin and eosin staining for review by an expert hepatobiliary pathologist (EEF), and the other half was disaggregated to generate single-cell suspensions for multiparameter FC.

**Figure 2.**
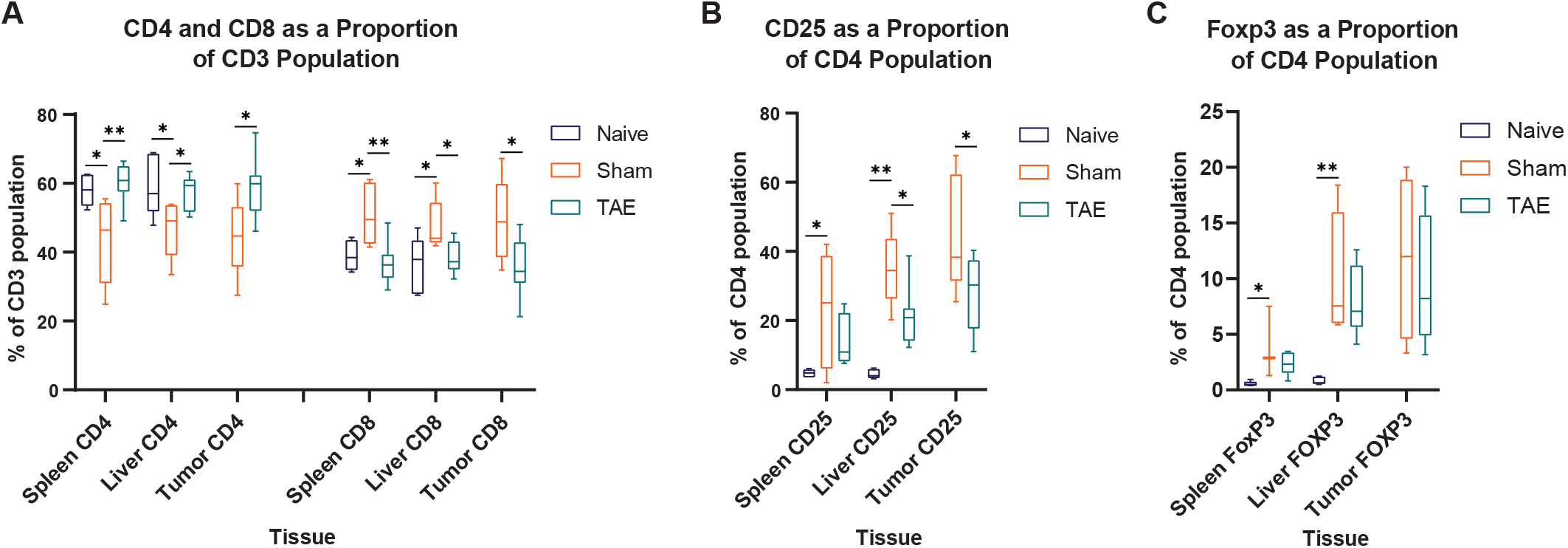
Cirrhosis and TAE Modulate Lymphocyte Populations. (A) Box and whisker plot showing the percentage of CD4+ and CD8+ cells as a proportion of the CD3+ population in spleen, liver, and tumor specimens from healthy DEN naïve rats that underwent sham embolization (Naïve, N=5 rats), cirrhotic DEN-treated rats that underwent sham embolization (Sham, N=5 rats), and cirrhotic DEN-treated rats that underwent conventional TAE with Embospheres (TAE, N=8 rats). (A) Box and whisker plot showing the percentage of CD25+ cells as a proportion of the CD4+ population. (B) Box and whisker plot showing the percentage of Foxp3+ cells as a proportion of the CD4+ population. \**p* < 0.05, \*\**p* < 0.01.

### TAE Enhances Recruitment of Tumor Infiltrating Lymphocytes

Next, in order to determine whether TAE results in an increased number of tumor infiltrating lymphocytes (TILs), we compared the average number of lymphocytes per mm^2^ within embolized and untreated tumors at post-embolization days 2, 7, and 12. Within embolized tumors there was a significantly greater number of CD3+, CD4+, and CD8+ TILs at 12 days post-embolization relative to untreated controls (191.4 vs 106.7, P<0.01; 127.8 vs 53.8, P<0.0001; 131.4 vs 78.3, P<0.01) (Figure 3a); however, no difference was observed in the average number of Foxp3+ TILs at any time point (Figure 3a). Since the proportion of TILs in stromal versus intratumoral compartments has been shown to be clinically significant in many types of cancer^35^, we also examined the average number and proportion of lymphocytes in both of these compartments. Consistent with the above results, we found a significantly greater number of CD3+, CD4+, and CD8+ cells within the intratumoral compartment at 12 days post-embolization relative to control tumors (123.9 vs 45.3, P<0.05; 80 vs 22.3, P<0.0001; 71.9 vs 35.9, P<0.001) (Figure 3b). Interestingly, we also found a significantly greater proportion of Foxp3+ cells within the intratumoral compartment at 2 days post-embolization (43.6% vs 21.0%, P<0.01) that did not persist at 7 or 12 days (Figure 3c).

**Figure 3.**
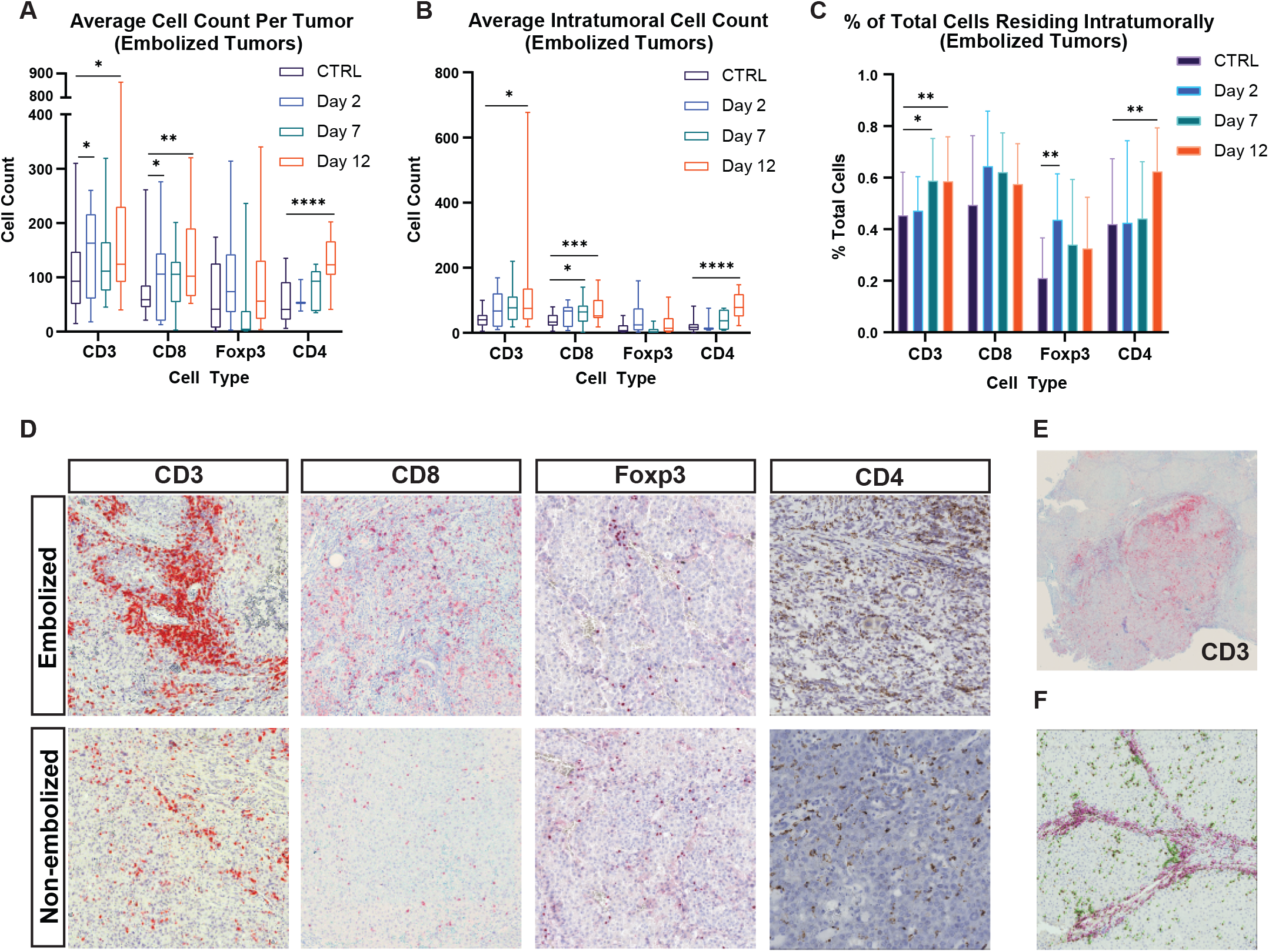
TAE Induces Increased Lymphocyte Infiltration in Embolized Tumors. (A) Box and whisker plot showing the average number of CD3+, CD8+, Foxp3+, and CD4+ cells at 2 days, 7 days, and 12 days post-embolization relative to control tumors that did not undergo embolization. (B) Box and whisker plot showing the average number of cells within the intratumoral compartment of embolized tumors at 2 days, 7 days, and 12 days post-embolization relative to control tumors that did not undergo embolization. (C) Bar chart showing the average percentage of cells found within the intratumoral compartment as a function of total number of cells (intrastromal + intratumoral cells) of embolized tumors at 2 days, 7 days, and 12 days post-embolization relative to control tumors that did not undergo embolization. (D) Representative images of histological staining of CD3, CD8, Foxp3, and CD4 cell markers in embolized and non-embolized tumors. (E) Low magnification image showing CD3+ cell infiltration into an HCC tumor 12 days post-embolization. (F) Example image demonstrating counting method used to quantify intrastromal cells (pink markers) from intratumoral cells (green markers) within HCC specimens. \**p* < 0.05, \*\**p* < 0.01, \*\*\**p* < 0.001, \*\*\*\**p* < 0.0001.

### TAE Increases the Number and Proportion of TILs in the Intratumoral Compartment of Non-Target Tumors

Since TAE has demonstrated the potential to induce distant tumor regression through the so called abscopal effect^36^, we sought to determine whether TAE influenced tumor lymphocyte infiltration in non-target HCC tumors in animals that were embolized. Although there was a trend toward increased numbers of TILs, no difference was observed in the average number of CD3+, CD4+, CD8+, and Foxp3+ TILs within the entire tumor at any of the time points studied (Figure 4a). However, non-target tumors demonstrated a significantly greater number of CD4+ and CD8+ TILs at day 12 relative to untreated controls (63.6 vs 22.3, P<0.05; 68.7 vs 35.9, P<0.05) (Figure 4b). Differences in the proportions of TILs within the intratumoral compartment of non-target lesions as compared to controls was heterogeneous, with a significantly greater proportion of intratumoral CD8+ TILS at day 7 (72.8% vs 49.4%, P<0.05), a significantly greater proportion of intratumoral CD4+ TILs at day 12 (58.1% vs 41.9%, P<0.05), and a significantly greater proportion of Foxp3+ TILs at days 7 and 12 (48.1% vs 21.0%, P<0.05; 43.6% vs 21.0%, P<0.01) (Figure 4c).

**Figure 4.**
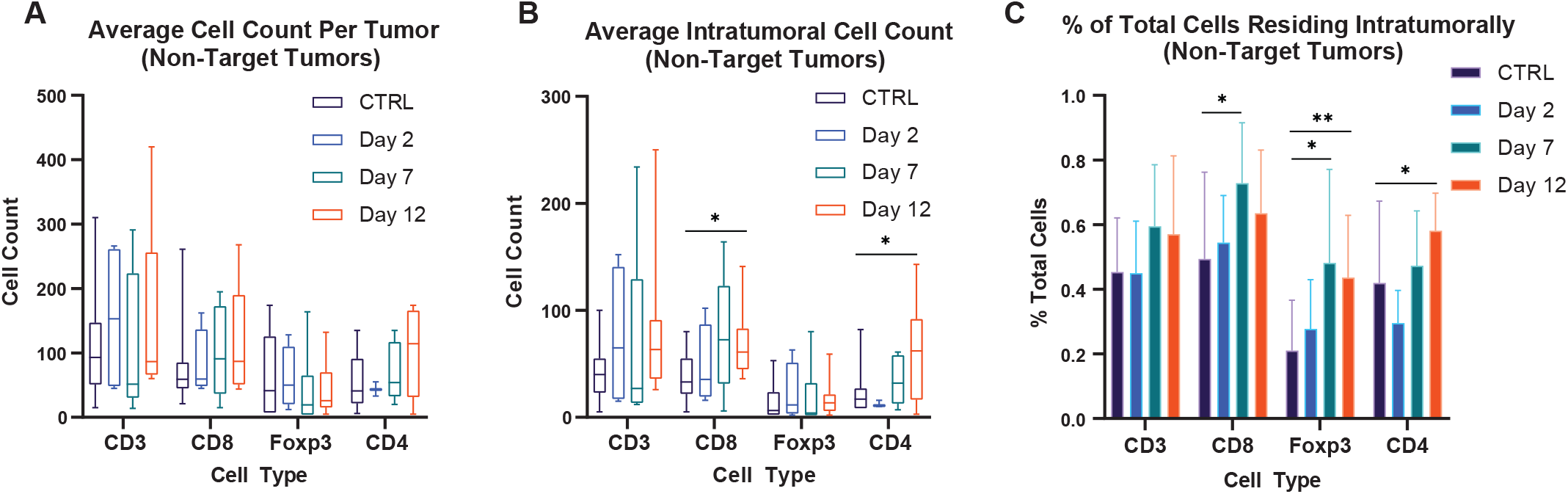
TAE Increases the Number and Proportion of TILs in the Intratumoral Compartment of Non-Target Tumors. (A) Box and whisker plot showing the average number of CD3+, CD8+, Foxp3+, and CD4+ cells at 2 days, 7 days, and 12 days post-embolization in non-target tumors relative to control tumors from rats that did not undergo embolization. (B) Box and whisker plot showing the average number of cells within the intratumoral compartment of non-target tumors at 2 days, 7 days, and 12 days post-embolization relative to control tumors. (C) Bar chart showing the average percentage of cells found within the intratumoral compartment as a function of total number of cells (intrastromal + intratumoral cells) in non-target tumors at 2 days, 7 days, and 12 days post-embolization relative to control tumors. \**p* < 0.05, \*\**p* < 0.01.

### TAE Induces Increased Expression of PD-L1 by HCC cells

PD-L1 tumor expression is considered a predictive biomarker of response to ICIs and is increased in HCCs^37–40^. Thus, we sought to determine whether TAE influences the expression of PD-L1 by HCC tumors in this model. With respect to control tumors, a greater than two-fold increase in the average expression level of PD-L1 was observed at 12 days post-embolization (4.1 vs 1.9, P<0.0001), as well as a smaller but significant increase in PD-L1 expression within non-target tumors (3.5 vs 1.9, P<0.05). (Figure 5).

**Figure 5.**
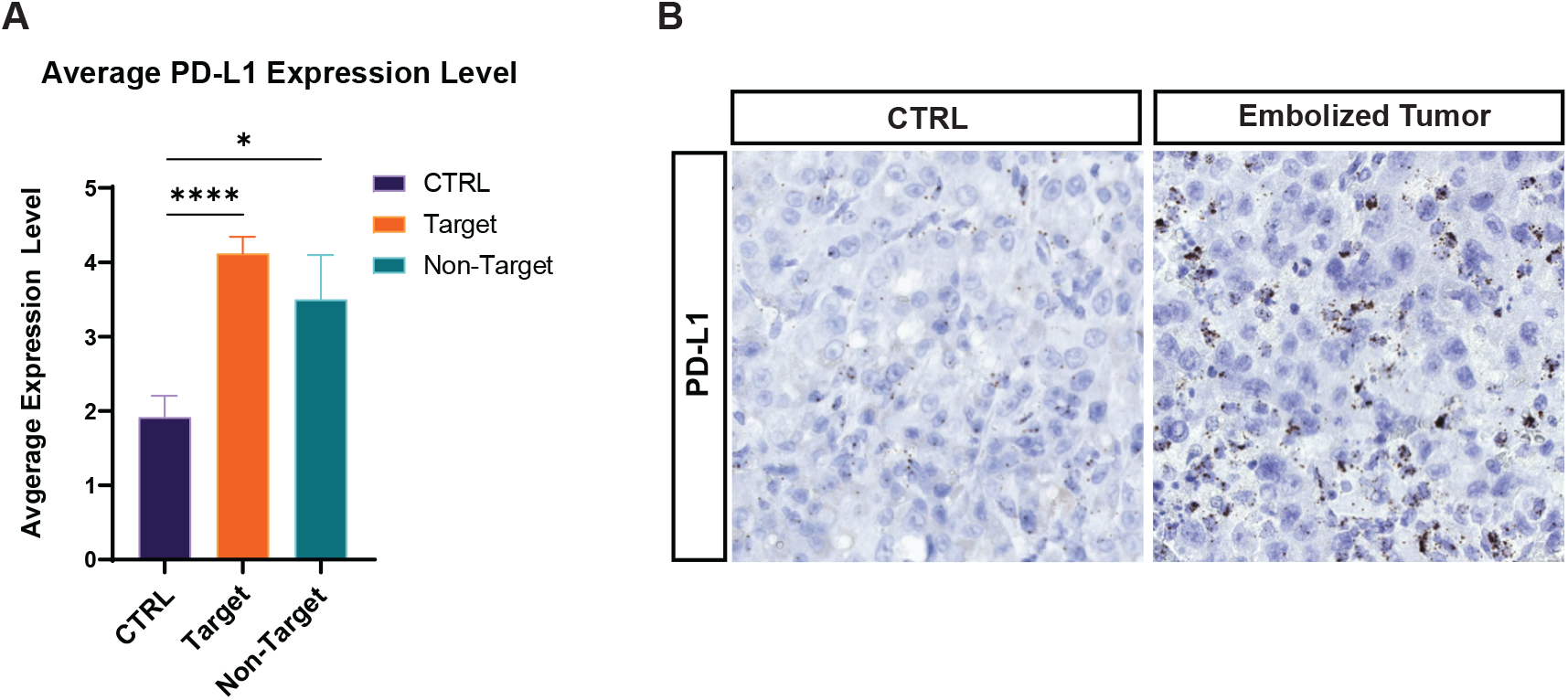
TAE Induces Increased Expression of PD-L1 by HCC cells. (A) Bar chart showing the average expression level of PD-L1 as determined with *in-situ* hybridization (ISH) in untreated control tumors, embolized tumors, and non-target tumors in rats that underwent embolization. (B) Representative images of PD-L1 ISH in embolized and untreated control tumors that did not undergo embolization. \**p* < 0.05, \*\*\*\**p* < 0.0001.

### Embolic Type and Location Influences Recruitment of Lymphocytes and Macrophages

In order to identify factors that might influence the immune response to TAE, we compared differences in the numbers of peri-bead lymphocytes and macrophages between embolic agents commonly used in the clinic: Embospheres (trisacryl gelatin) and LUMI beads (polyvinyl alcohol). H&E stained sections demonstrated a strikingly greater number of inflammatory immune cells encircling Embospheres compared to LUMI beads (Figure 6a). These findings were confirmed on quantitative analysis of IHC which demonstrated a significantly greater number of CD3+, CD4+, and CD8+ lymphocytes in proximity to Embospheres as compared to LUMI beads (4.1 vs 2.0, P<0.01; 3.7 vs 2.0, P<0.05; 2.2 vs 1.1, P<0.05) (Figure 6b). There was no difference in FoxP3+ cell numbers (1.2 vs 1.2, P>0.05) (Figure 6b). Regression analysis revealed a linear relationship between the number of embolic particles and the number of leukocytes adjacent to embolic. Similarly, IHC demonstrated a markedly greater number of CD68+ cells around Embospheres (13.6 vs 5.8, P<0.0001), with a surprising decrease in the number of CD163+ cells (inhibitory subset of CD68+ cells) as compared to LUMI beads (0.8 vs 1.9, P<0.05) (Figure 6c). In addition, the observed Embosphere-associated recruitment of CD3+, CD4+, CD8+, FoxP3+, and CD68+ cells increased with time following TAE (1.8 vs 4.9, P<0.01; 1.0 vs 5.0, P<0.0001; 1.0 vs 2.0, P<0.05; 0.2 vs 1.8, P<0.0001; 10.8 vs 16.6; P<0.05) (Figure 6C).

**Figure 6.**
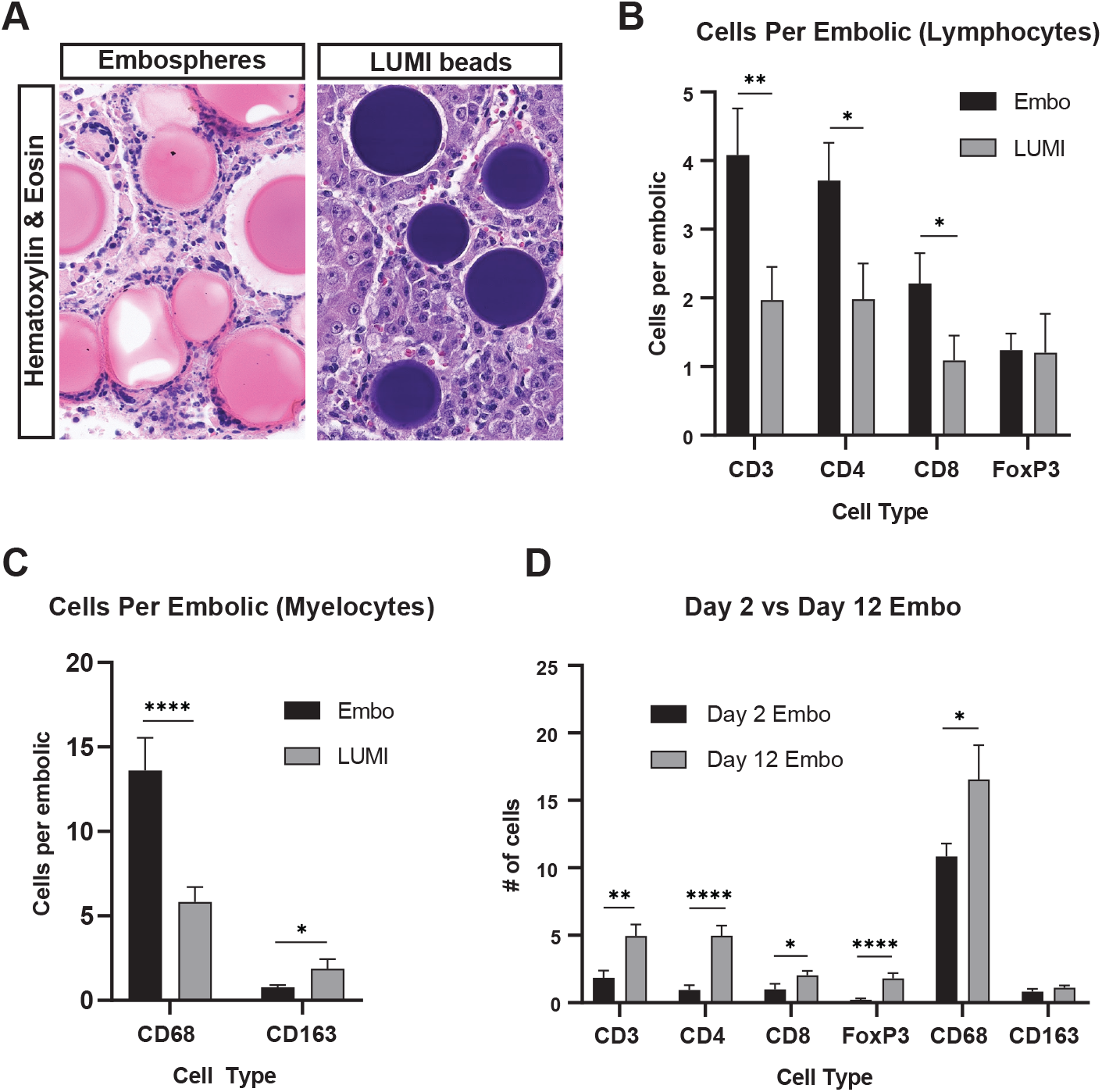
Differential Recruitment of Lymphocytes and Myelocytes Based on Type of Embolic. (A). Representative images of hematoxylin and eosin staining demonstrating differences in foreign body reactions induced within HCC tumors embolized with LUMI Beads or embospheres. (B). Bar chart showing the average number of CD3+, CD4+, CD8+, and Foxp3+ cells per embolic by embolic type. (C). Bar chart showing the average number of CD68+ and CD163+ macrophages per embolic by embolic type. (D). Bar chart showing the average number of cells surrounding Embospheres at 2 days and 12 days post-embolization. \**p* < 0.05, \*\**p* < 0.01, \*\*\*\**p* 0.0001.

To compare the influence of intra-vs. extravascular location of the embolic on macrophage recruitment, we performed IF for the general macrophage marker CD68 as well as the vascular marker SMA (Figure 7a). Quantitative analysis revealed that Embospheres exited vessels at significantly higher rates as compared to LUMI beads (87.4% vs 41.2%, P<0.001) (Figure 7b). Interestingly, embolics which have extravasated recruit a greater number of CD68+ macrophages as compared to embolics that remain within the intravascular space (Figure 7a, 7c). Indeed, both Embospheres and LUMI beads demonstrated a larger peri-embolic CD68 signal when outside of the vessel as compared to within (17.9 vs 7.0, P<0.0001 for Embospheres, 6.4 vs 3.4, P<0.05 for LUMI beads) (Figure 7a, 7c).

**Figure 7.**
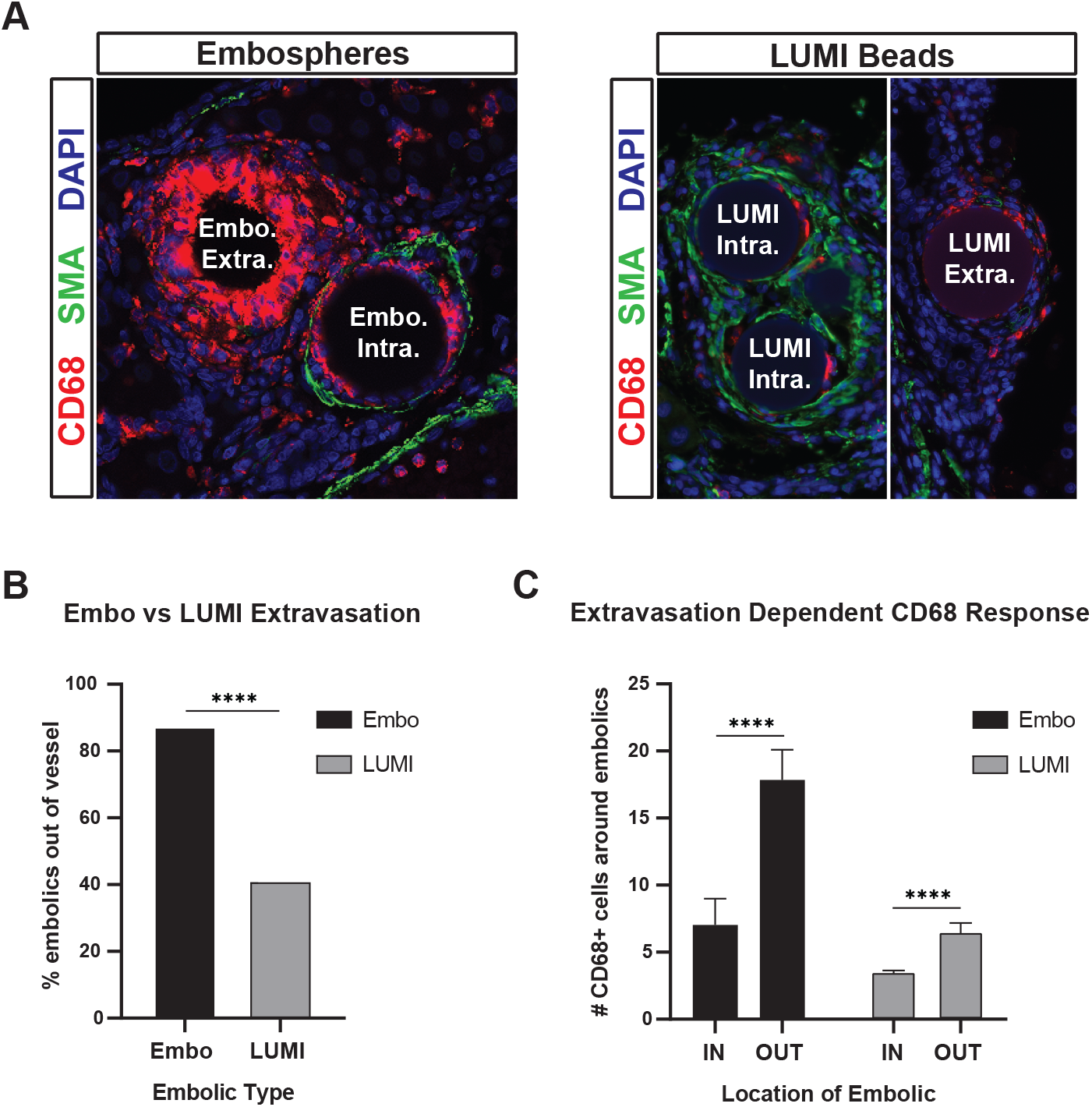
Embospheres Induce a Marked Foreign Body Reaction that Increases with Extravasation. (A). Representative images showing triple immunofluorescence for the macrophage marker CD68, arterial marker smooth muscle actin (SMA), and DAPI on HCC specimens containing LUMI Beads and Embospheres. The foreign body reaction, as a measure of macrophage number, differs between embolic type and location. (B). Bar chart showing the percentage of embolics found within SMA+ vascular walls by bead type. (C). Bar chart showing the average number of CD68+ macrophages surrounding LUMI Beads and Embospheres based upon vascular location. Abbreviations: Embo (Embospheres); Extra (extravascular); Intra (intravascular). \*\*\*\**p* < 0.0001.

### Human Data

Pending

## Discussion

The incidence of HCC is increasing across the globe^41^. Within the United States, HCC ranked number one in terms of increased cancer mortality between 1990 and 2015^42,43^. These dismal statistics issue, at least in part, from the fact that there remain limited therapeutic options for patients with HCC. Given the potential for locoregional therapies to engage the immune system, and the well documented connection between patient outcomes and immune response, many clinical trials are already underway which are designed to test the presumed benefit of combination locoregional and immunotherapy^44^; however, there remains a paucity of data characterizing the influence of embolotherapy on the immune landscape that is required to inform these trials.

In addition to the unique immunotolerant features of the liver^45–47^, HCC has the demonstrated ability to evade both endogenous immune responses and engineered immunotherapies through multiple mechanisms including increased frequencies of regulatory T cells and myeloid-derived suppressor cells, impairment of tumor antigen processing and presentation, alterations in the tumor metabolic environment which are hostile to immune function, as well as upregulation and overexpression of inhibitory receptors such as immune checkpoint inhibitors including programmed death ligand 1 (PD1)^18,48,49^. Thus, while infiltrating immune cells have the potential to halt tumor growth, they can also help to create an immunosuppressive environment that facilitates tumor progression^50^. This complex interplay between immune cells, cancer cells, and supporting stromal cells has been the topic of numerous studies aimed at identifying prognostic markers and therapeutic targets in the fight against HCC. While the role of various immune cell subtypes in anti-tumor immunity is still somewhat controversial, it is generally accepted that increased numbers of CD8+ cytotoxic T-cells predict prolonged survival whereas macrophages, mast cells, and Foxp3+ T-regulatory cells are associated with poorer outcomes^50,40,49,51^.

Here, we show that TAE has significant effects on the number and ratio of effector T cells within the tumor microenvironment, and that the expression of PD-L1 is in increased in both target and non-target tumors. We further examined the impact of embolic type and location on immune cell recruitment, demonstrating striking differences in the resulting immunomodulation. Lastly, we corroborated our findings in a cohort of human HCC samples, emphasizing the implications of these results for informing future translational studies.

Consistent with the findings of the current data, prior studies that have examined the impact of TAE on tumor infiltrating lymphocytes have demonstrated significant alterations in the tumor immune microenvironment. Using a rabbit VX2 liver tumor model, Duan et al found that following TAE, the number of CD8+ T cells surrounding embolized tumors was increased relative to control^52^. Our data not only support this finding, but expand upon it by integrating data highlighting the importance of cellular localization within tumors to either the intratumoral or intrastromal compartment. In addition, Avritscher et al found no difference in the percentage of Foxp3, IFNγ-producing CD4, or CD8+ T-cells within tumor, tumor margin, or adjacent non-cancerous liver after embolization using an orthotopic rat model of TA(C)E and HCC^53^. While Avritscher et al found no difference in the percentage of immune cells, they did not examine total numbers of intrahepatic immune cells, underscoring the challenges of integrating flow cytometry data with immunohistochemistry. The current study applied both of these methodologies enabling the identification of decreases in the ratio of certain immune cell populations (e.g. CD8+ T cells) consistent with the prior study, with a concomitant increase in the overall number of CD8+ T-cells following TAE. Lastly, Craciun et al retrospectively examined resection specimens from patients undergoing partial hepatectomy for HCC with preoperative TACE, and found no difference in the expression of CD3, CD4, or CD8 when compared to controls^54^. Such discrepancy may be in part due to differing patient populations and methodologies; for example, the use of gelfoam as the embolic agent in a portion of the patients and the collection of samples at 14-15 weeks following TACE. Here, we demonstrate that cell type specific immune responses evolve over time and differences in embolics may dramatically alter induced changes in the immune microenvironment.

Indeed, the choice of embolics varies widely across TA(C)E protocols emphasizing the importance of exploring the role of embolics in immune cell recruitment. H&E staining revealed dramatically different foreign body responses in HCCs treated with LUMI beads as compared to Embospheres. Further analysis demonstrated significant differences in the quantity of lymphocytes and myelocytes attracted to each type of embolic. We also showed a time dependent accumulation of cells around Embospheres, with significant increases in most cell types from days 2 to 12, indicating a progression of the immune response over time. Interestingly, we also found that Embospheres were more than twice as likely to extravasate outside of the vessel wall and into the tumor parenchyma compared to LUMI beads. Embospheres located extravascularly recruited over twice as many CD68+ macrophages than Embospheres that remained within vessels. These results demonstrate that there are significant differences between embolic particles in terms of how they access a tumor and engage the immune system, with intravascular embolics having a limited ability to modulate the immune system. Extravasation may therefore be an advantageous phenomenon that techniques such as pressure-enhanced delivery may be able to harness. Such findings may also have important implications for the design of embolic particles engineered to induce specific immune responses. To corroborate the clinical significance of our findings, we performed similar measures of immune cell recruitment on a cohort of human HCC specimens following TACE (PENDING).

Beyond its effect on target tumors, the described data demonstrate that TAE alters the immunobiology of distant, non-target tumors as demonstrated by an increased number of CD4, CD8, and Foxp3+ cells in the intratumoral compartment. This finding holds important implications given the growing body of literature demonstrating that the prognostic value of TILs is relative to their distribution within peritumoral, intratumoral, or stromal compartments^35^. As such, the finding of an increased number of T cells within the intratumoral compartment of non-target tumors reaffirms the potential for a TAE-induced abscopal effect resulting from the release of novel tumor associated antigens and recruitment of effector T cells. This potential is further supported by the significant alterations in T-cell populations identified in the spleens of TAE-treated rats.

These findings hold important implications given the completed^22,23^ and ongoing clinical trials investigating the use of TACE in combination with immune checkpoint inhibitors (ICIs) for the treatment of HCC (for a complete list of trials, visit clinicaltrials.gov). In addition to demonstrating the temporal changes in the immune response induced by TACE, we found that TAE causes increased expression of PD-L1 on both target and non-target tumors. These data reaffirm the notion that combination therapy with PD-1 inhibitors may act synergistically to overcome the seemingly negative consequence of increasing PD-L1 expression on tumor cells, which might otherwise promote T-cell exhaustion. ICIs have also been associated with immune related adverse reactions that can sometimes be fatal. A thorough understanding of the immune response to TA(C)E will likely have important implications for informing the administration of ICIs, both to optimize treatment response and to minimize adverse reactions. In addition, catheter directed local delivery of ICIs directly into tumors has further potential to reduce systemic adverse effects while allowing for increased intratumoral effective concentration. Future studies examining markers of immune cell activation and exhaustion in the setting of immune checkpoint blockade will help to further elucidate the significance of our findings. These functional assessments will be crucial in determining which ICIs can most effectively engage the immune system as well as the optimal peri-procedural timing of their administration.

Finally, the described data underscore the importance of animal models that faithfully recapitulate the biology of human disease. DEN-induced autochthonous HCCs develop in a background of hepatic cirrhosis which results from hepatocellular injury and is known to play an important role in the pathogenesis of most human HCCs in the US^55,56^. Consistent with human HCC, we show that cirrhosis significantly alters the local immune microenvironment and that TAE induces changes that mitigate these alterations. Moreover, autochthonous tumors have been shown to most closely recapitulate the biology of the tumor microenvironment, including vasculature, immunobiology and architecture^57–59^. Specifically, the dependence of HCC on the arterial blood supply develops over time through the growth of abnormal intranodular arteries, a characteristic that has been demonstrated to distinguish autochthonous from transplanted tumor models and is particularly relevant in the study of embolotherapy^60,61^. Indeed, the observed transit of embolics out of the vasculature and into the tumor interstitium is likely to be influenced by the unique structural features of intratumoral arteries. The demonstration of this phenomenon in clinical samples further emphasizes the translational relevance of this model. In addition to translational relevance, the DEN-induced rat model of HCC and TAE provides a feasible and cost-effective solution to the study of TAE-induced alterations based on the low procurement and husbandry fees, ease of handling, and availability of commercial reagents. While models in other species offer potential advantages, the high costs and dearth of reagents for these models, such as species specific antibodies for immunohistochemistry or flow cytometry, pose significant limitaitons^62^.

It is important to recognize the limitations of our study. Although we have attempted to characterize the various cellular alterations which follow TAE, we have yet to determine functional changes that occur in terms of immune cell activation and exhaustion. We have also focused primarily on lymphocytic cells, but recognize that a variety of myelocytic cell types not addressed here play critical roles within the tumor microenvironment. In addition, due to the nature in which our samples were collected, quantitative measures of immune cell number were not obtained using stereological methods, potentially biasing our results. Furthermore, we only examined the impact of bland embolization on the immune response, and thus it is unknown how the addition of chemotherapeutic agents might affect our results.

Taken together, this study represents the first comprehensive report on cellular immune responses in HCC following TAE within an autochthonous animal model. These data demonstrate the potential of locoregional embolotherapy to modulate the tumor immune microenvironment directly and provide unique insights into the nature, breadth and mechanism of the induced alterations. These findings hold important implications for the on-going development of novel therapeutic strategies combining locoregional therapy with immunomodulators as well as for the development of techniques and materials that can further leverage the unique modulation of the tumor immune microenvironment that can be achieved through locoregional endovascular therapies.

## Materials and Methods

### Autochthonous rat model of HCC and TAE

Animal studies were conducted according to institutionally approved protocols for the safe and humane treatment of animals. Briefly, autochthonous HCCs were induced in male Wistar rats (Charles River Laboratories) using an established protocol including ad libitum oral intake of 0.01% diethylnitrosamine (DEN) for 12 weeks^63^. Rats with tumors 0.5-1.0 cm in maximal diameter on T2-weighted MRI were selected for embolization. Arterial access was gained through ventral tail artery approach. Using fluoroscopic guidance, arteries feeding the tumor underwent catheterization and embolization with 0.2 mL of 40–120 μm Embosphere particles (Merit Medical, South Jordan, Utah) or 70–150 μm LUMI™ beads (BTG, London, UK) suspended in 1 mL iopamidol contrast medium (Isovue 370; Bracco, Monroe Township, New Jersey) was performed as previously described^63^. Pre- and post-embolization arteriography was performed using an AngioStar Plus Imaging System (Siemens, Malvern, Pennsylvania) in order to confirm occlusion.

### Tissue preparation and histology

After the desired number of post-treatment days, treated and untreated tumors were harvested from rats according to institutionally approved protocols following euthanasia. Tissue was fixed in formalin and dehydrated with 70% alcohol prior to paraffinization. The percentage of viable and necrotic HCC tissue was estimated from hematoxylin and eosin (H&E) stained sections by a hepatobiliary pathologist with extensive experience in evaluating HCC (E.E.F).

### Immunohistochemistry, Immunofluorescence and microscopy

Standard immunohistochemistry (IHC) and Immunofluorescence (IF) protocols were used to stain formalin-fixed, paraffin embedded tissues. Briefly, after deparaffinization and rehydration of 4-μm-thick tissue sections, heat induced epitope retrieval was performed in 10 mM sodium citrate buffer (pH 6.0) using a pressure cooker for 20 min. The slides were then blocked for 30 min in StartingBlock™ T20. Primary antibodies used in this study included: anti-CD8a (eBioscience, OX8, 1:200), anti-CD3 (Abcam, ab5690, 1:200), anti-Foxp3 (eBioscience, FJK-16s, 1:200), anti-CD68 (BioRad, ED1, 1:200), anti-CD163 (BioRad, ED2, 1:200), and anti-alpha smooth muscle actin (Abcam, ab5694, 1:1000). Primary antibodies were diluted in 5% bovine serum albumin diluted in PBS with 0.2% Triton-X and incubated with sections overnight at 4°C. Secondary antibodies of the appropriate species conjugated to either alkaline phosphatase or horseradish peroxidase (Vector Labs) were then applied for 30 min at room temperature. Color was developed according to manufacturer protocols using DAB (brown) or Vector Red (red) substrate kits and counterstained with Mayer’s hematoxylin. For IF, secondary antibodies were conjugated to Alexa fluorophores (488 or 568, Invitrogen). DAPI (300nM) was applied concurrently with secondary antibodies to label cell nuclei. Entire tissue sections were then imaged with a 20x objective using a BZX fluorescence microscope (Keyence, Osaka, Japan) with Keyence imaging software.

### In-situ hybridization (ISH)

ISH for CD4 and PDL1 was performed on FFPE sections using RNAscope 2.5 HD Brown Reagent Kit in combination with RNAscope Probe-Rn-CD4 and RNAscope Probe-NPR-0002409-PDL1 (Advanced Cell Diagnostics, Inc., Hayward, CA, USA) according to the manufacturer’s instructions.

### Flow Cytometry

Rat tissue was prepared for flow cytometry based on established protocols^64^. Briefly, once the appropriate time point was reached, rats were euthanized and the portal vein was perfused with cold PBS. Tissues were collected and kept in cold PBS with 2% fetal bovine serum (FBS). Tissues were strained through a 70um filter and washed twice with 2% FBS at 500 × g for 5 min at 4°C. Immune cells were separated from hepatocytes via gradient centrifugations using 33.75% Percoll (GE Healthcare) in PBS at 700 × g for 12 minutes at room temperature. Isolated leukocytes were carefully isolated and washed with 2% FBS at 340 × g for 5 min at 4°C. RBC lysis was performed with red blood cell lysis buffer (Sigma-Aldrich) for 4 min. Cell suspension was then mixed with heat inactivated FBS and centrifuged at 340g × for 5 min at 4°C.

Cells were then counted, washed in PBS, and stained with a Near-IR Live-Dead stain (Thermo Fisher, 2:1000) at 10×6 cells per 1 ml for 20 minutes. Samples were washed twice with flow buffer (Fisher Scientific) at 500 × g for 5 min at 4°C and blocked with mouse serum (1:20 in flow buffer) for 10 minutes. Cells were then spun down at 500 × g for 5 min at 4°C and resuspended with the following antibody-conjugated fluorophores for 20 minutes at room temperature: anti-CD3-BV421 (BD, 1F4, 1:100), anti-CD4-PE-Cy7 (BD, OX-35, 1:100), anti-CD8-BV510 (BD, OX8, 1:100), anti-CD25-APC (Thermo Fisher, OX39, 1:100), and anti-CD45-PE-Cy5 (BD, OX1, 1:100). After surface staining, cells were fixed, washed twice with flow cytometry buffer at 500 × g for 5 min at 4°C, permeabilized using a fix perm kit (Life Technologies), and stained with either anti-FoxP3-FITC (Thermo Fisher, FJK-16s, 1:100) or the isotype control (Invitrogen, eBR2a, 1:100) for 30 minutes at room temperature. FMO control for anti-CD25-APC was included. Cells were acquired using an LSRII flow cytometer (Beckman Coulter) and data was analyzed with FlowJo (Tree Star Inc).

### Cell Counting and Image Analysis

Cell counting was performed using ImageJ^65^ in a blinded fashion. Briefly, stromal and intratumoral compartments were defined according to standardized methods^35^. Tissue sections were screened at medium power (x100) in order to identify areas for analysis that would ensure representativeness and homogeneity. A quantitative evaluation of lymphocytes or macrophages was then performed at high power (x200) by counting the total number of cells within three separate areas each measuring 1 mm^2^. The average number of cells per mm^2^ was then calculated. The stromal and intratumoral compartments were both analyzed separately and cells per mm^2^ was reported. In order to obtain the percentage of cells within the intratumoral compartment, the average number of cells per mm^2^ within the intratumoral compartment was divided by the sum of the average number of cells per mm^2^ in both compartments. For CD4 quantification, a cell was considered positive if it had morphological features of a lymphocyte and contained at least 6 individual puncta or dense chromogen staining.

For quantification of PDL1 mRNA expression, tissue sections were screened as above in order to identify areas for analysis that would ensure representativeness and homogeneity. PD-L1 staining intensity within tumor cells was classified into 5 levels. Level 0, no expression; level 1: 0-10% of cells containing at least 2 puncta; level 2, 10-40% of cells containing at least 2 puncta; level 3, 40-70% of cells containing at least 2 puncta; level 4, 70-100% of cells containing at least 2 puncta.

For bead analyses, samples that had visible embolic particles were selected. In order to calculate peri-bead leukocytes, we created a region of interest (ROI) calculation which was twice the size of the bead’s diameter. When ROIs of several embolics were adherent, this was considered to be a single ‘island’ of embolics and all adjacent cells were counted together along with the number of embolic particles present. Embolic particle islands analyzed were capped at 50 visible particles. To calculate CD68 response while in or out of the vessel, we used the same ROI methodology and noted positive SMA staining around embolic particles as a way to signify intravascular positioning.

### Statistical analyses

Due to repeated measures being taken for many of the rats, Generalized Estimating Equation (GEE) Models were employed to determine the impact of the predictors (day, bead type, group) on the various outcome measures. A cutoff of .05 was used to determine statistical significance. Within graphs, error bars on box and whisker plots indicate 5-95^th^ percentile. Error bars on bar chart indicate standard error of the mean (SEM).

## Data availability

The datasets generated during the current study are available from the corresponding author on reasonable request.

## Acknowledgments

VA Merit I01-CX001933 (DEK,TPG)

## Notes

### Competing Interest Statement

The authors have declared no competing interest.

